# An *Agrobacterium* strain auxotrophic for methionine is useful for switchgrass transformation

**DOI:** 10.1101/2022.02.24.481806

**Authors:** Mónica Prías-Blanco, Timothy M. Chappell, Emily F. Freed, Eudald Illa-Berenguer, Carrie A. Eckert, Wayne A. Parrott

## Abstract

Auxotrophic strains of *Agrobacterium tumefaciens* can contribute to the development of more efficient transformation systems, especially for crops historically considered recalcitrant. Homologous recombination was used to derive methionine auxotrophs of two common *A. tumefaciens* strains, LBA4404 and EHA105. The EHA105 strains were more efficient for switchgrass transformation, while both the EHA105 and LBA4404 strains worked equally well for the rice control. Event quality, as measured by transgene copy number, was not affected by auxotrophy, but was higher for the LBA4404 strains than the EHA105 strains. Ultimately, the use of auxotrophs reduced bacterial overgrowth during co-cultivation, with a concomitant decrease in the need for antibiotics. In turn, reduced overgrowth allowed longer co-cultivation periods, with a trend towards higher transformation frequencies for switchgrass, but not for rice.

## Introduction

The bacterium, *Agrobacterium tumefaciens*, revolutionized plant science over the past decades due to its ability to transfer DNA into plant cells. This success is in large part due to its ability to infect and transform a wide host-range of plant species in the laboratory (Nester, 2015). Yet, despite *Agrobacterium*’s popularity, the strains available within the public sector lack modifications that could simplify procedures or increase overall plant transformation efficiency. Successful transformation depends on the ability to control *Agrobacterium* using antibiotics to prevent it from overgrowing the explants, but antibiotics can have undesired effects on the plant tissue (Shackelford and Chlan, 1996). Despite the usefulness of using antibiotics to control agrobacteria in tissue culture, control is not always successful, and some antibiotics can affect tissue growth (Meng et al., 2014).

To reduce the need for using antibiotics, Collens et al. (2004) were the first to develop and use auxotrophic disarmed strains of *Agrobacterium* explicitly for transformation. They used transposon mutagenesis to develop auxotrophs of EHA105 (Hood et al., 1993) focusing on mutants for adenine, leucine, and cysteine that had no growth or reduced growth in the absence of the required nutrient. When these auxotrophic mutants were tested on *Nicotiana glutinosa* cells, they exhibited higher rates of transformation as measured by GUS staining and a quantitative GUS assay than wild type EHA105. In particular, the cysteine auxotroph showed a GUS expression level 85-fold higher than the corresponding prototrophic strain. However, transformation frequency based on GUS expression when using the parental EHA105 was close to zero. This lower-than-expected transformation frequency with the wild-type strain inflates the efficiency of the auxotroph. Collens et al. (2007) and Larsen and Curtis (2012) later showed the utility of the cysteine auxotroph for stable transformation to evaluate viral vector components.

In the private sector, an auxotrophic strain for methionine derived from LBA4404 (Hoekema et al., 1983) was developed though X-ray irradiation. To further increase transformation efficiency, additional *vir* genes from *A. tumefaciens* A281 were added by placing them on a ternary plasmid, pTOK47, to improve virulence (Jin et al., 1987). The strain, referred to as LBA4404metHV, was patented for dicot transformation. Similarly, the strain ‘ATHVade, his’, which was derived from a UV-light mutagenized AGL0 strain (Lazo et al., 1991) and is a double adenine and histidine auxotroph, (Dirks and Peeters, 2001) was also patented. Importantly, auxotrophy did not affect the transformation efficiency of the strains when tested on *N. tabacum*. The ‘ATHVade, his’ strain exhibited a 224% transformation efficiency while its parental AGL0 yielded 230% based on the number of transformed shoots per infected leaf segment.

More recently, a LBA4404 strain auxotrophic for thymidine (Ranch et al., 2010) was used for stable transformation of maize and other monocots (Anand et al., 2018; Hoerster et al., 2020; Lowe et al., 2018; Lowe et al., 2016). The LBA4404Thy-strain was generated using homologous recombination, and when combined with the superbinary vector pSB1 and the binary plasmid PHP15303, resulted in a 23% transformation frequency in maize while the same plasmid combination with the parental strain showed 26% using the wild-type strain with the same plasmid combination.

Despite the potential benefits of using auxotrophic strains of *Agrobacterium*, such strains are generally not available in the public sector. Towards that end, Aliu et al. (2020) used homologous recombination to develop thymidine auxotrophs of strains EHA101, EHA105, and EHA105D. Transient GUS expression assays using hyper-transformable seedlings (Wu et al., 2014) of arabidopsis showed that, except for EHA105D, auxotrophic strains can deliver T-DNA at comparable levels to the parental strains. In addition to being a thymidine auxotroph, EHA105D has a deleted *atsD* gene, which is implicated in plant cell attachment. However, the LBA4404Thy-auxotroph remained the most effective strain (Aliu et al., 2020).

This work was started to develop additional auxotrophic strains of *Agrobacterium* and make them available in the public sector. Switchgrass transformation was chosen to validate these strains by evaluating their performance in a difficult transformation system. Rice transformation was used for comparison. Since a common challenge when using *Agrobacterium* lies in controlling the overgrowth of *Agrobacterium*, here we show auxotrophs of EHA105 and LBA4404 maintain T-DNA transfer ability without overgrowing the tissue explants.

## Methodology

### Plasmid and *Agrobacterium* strains

Auxotrophs for methionine were generated for *Agrobacterium tumefaciens* EHA105 and LBA4404 using a protocol developed for constructing *Pseudomonas syringae* pv. Tomato DC3000 mutants with minor modifications (Kvitko and Collmer, 2011). The Gateway-compatible vector pDONR1K18ms (Addgene plasmid #72644) was used to construct the plasmids. As deleting bacterial genes relies on homology to flanking DNA, PCR primers (Table S1) were used to amplify flanks of the homoserine O-succinyltransferase (*metA*) gene (Table S2), the first enzyme in the methionine biosynthesis pathway in *Agrobacterium tumefaciens* (Rotem et al., 2013). PCR was performed using Q5™ High-Fidelity 2x Mastermix (New England Biolabs Inc., Ipswitch, MA, USA) with 40 ng genomic DNA (extracted with the DNeasy Blood and Tissue Kit, Qiagen, Valencia, CA, USA) and primers at 0.5 µM. PCR conditions were initial denaturation at 98 °C for 1 min, 35 cycles of (1) denaturation at 98 °C for 10 sec, (2) annealing temperature for 30 sec, and (3) extension at 72 °C for 30 sec, and final extension at 72 °C for 5 min.

Amplified flanks were gel-purified with the QIAquick Gel Extraction Kit (Qiagen, Valencia, CA, USA) and eluted in water before being joined using overlap extension PCR. The final sequence with the combined flanks was gel purified and eluted in water before being recombined into pDONR1K18ms using BP Clonase™ II (Invitrogen, Waltham, WA, US). The product from the BP Clonase II reaction was transformed into *E. coli* DH5a cells using standard heat shock methods and plated onto solid LB (Luria-Bertani) medium supplemented with 50 mg L^-1^ of kanamycin. After confirming the plasmid sequence to be correct, the deletion constructs (GenBank accessions OK181160 and OK181161) were electroporated into competent cells of EHA105 or LBA4404 as specified in the manual of the BioRad MicroPulser™ (Biorad Laboratories, Hercules CA, US), and the cells were allowed to recover at 28 °C for 3 hours for EHA105 or overnight for LBA4404 in liquid YM (Yeast Mannitol) medium without antibiotics.

After recovery, cultures were plated onto solid YM medium containing kanamycin at 50 mg L^-1^. The medium for EHA105 strains also had 50 mg L^-1^ rifampicin. Importantly, these plates were supplemented with an amino acid dropout mix, DO supplement -Ura (Takara Bio USA, Inc. Mountain View CA, US) to ensure adequate amino acid availability during the deletion process. YM medium was solidified with 1.5% w/v Bacto™ Agar (BD Diagnostics, Franklin Lakes, NJ, USA). Then two to eight individual colonies were picked and restruck on the above described medium to ensure the presence of the deletion plasmid. Then, cells from each restruck colony were resuspended in 1.5 mL of liquid YM medium and plated onto solid YM medium plates with 7.5% sucrose, in addition to the antibiotics and DO supplement.

The resulting colonies were screened for the targeted deletion using standard PCR using primers at 0.25 µM (Figure 1, Table S1) and OneTaq™ Polymerase (New England Biolabs Inc., Ipswitch, MA, USA) PCR conditions were initial denaturation at 95 °C for 7 min, 35 cycles of (1) denaturation at 95 °C for 30 sec, (2) annealing temperature for 30 sec, and (3) extension at 68 °C for 2 min, and final extension at 68 °C for 10 min. Confirmed colonies were streaked on solid YM medium containing the appropriate antibiotics and DO supplement. To confirm auxotrophy, the mutants were tested for growth in M9 minimal medium with or without the DO supplement. Confirmed strains that displayed growth in M9 minimal medium with the DO supplement but did not grow in M9 medium lacking the DO supplement were used in the plant transformation experiments.

**Figure 1.**
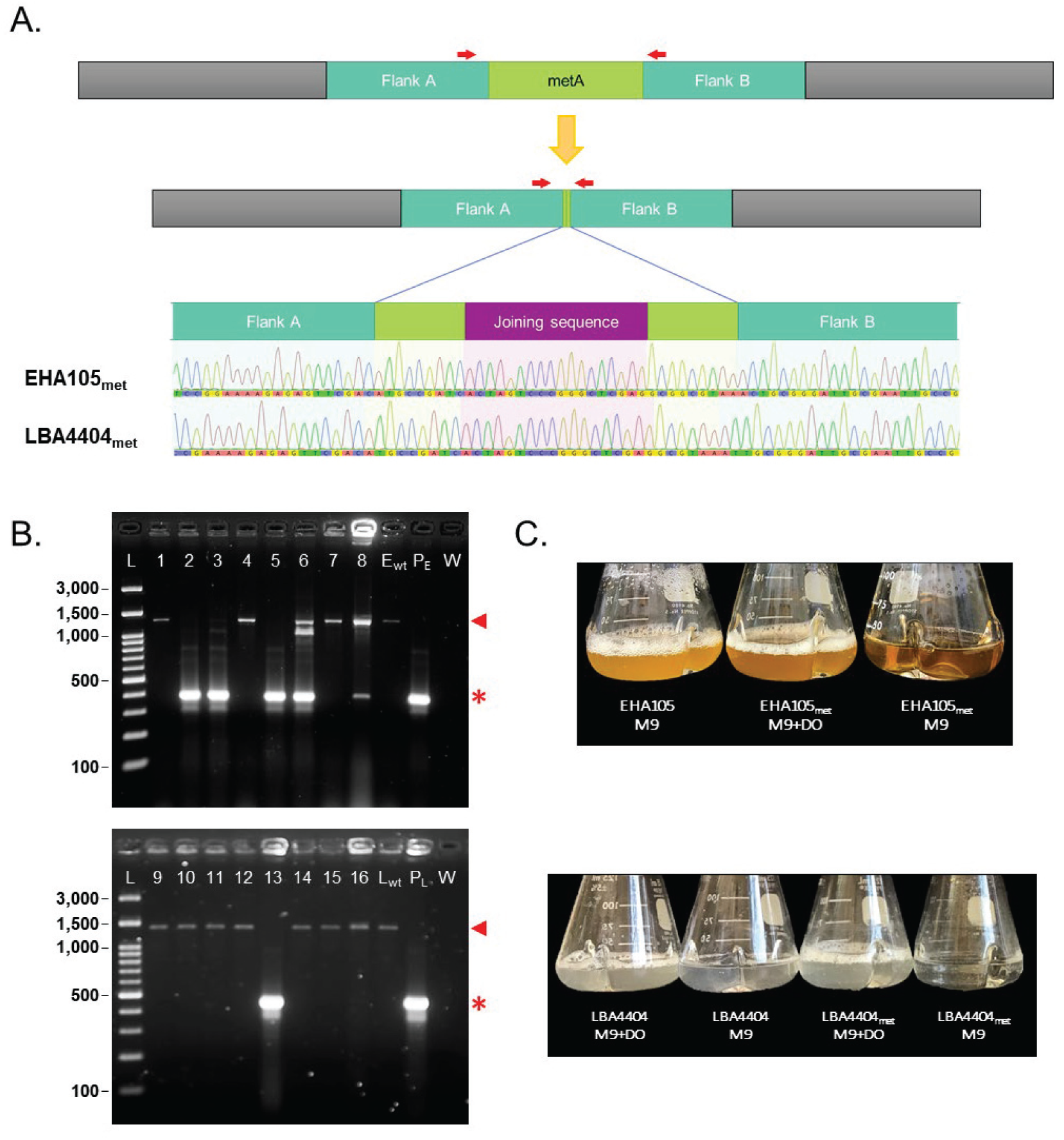
Generation of homologous recombination-mediated methionine auxotrophic mutants in *Agrobacterium*. (A) To make an auxotrophic *Agrobacterium* mutant we deleted the homoserine O-succinyltransferase (*metA*) gene – leaving the first three codons followed by the joining sequence CTCGAGCCCGGGACTAGT and the last two/three codons, including the stop codon of *met*A gene. Red arrows indicate the relative positions of the PCR primers used to validate the *met*A deletion. (B) Screening for *Agrobacterium metA* knockout mutants. Lanes 1 to 8: EHA105 background colony screening; E_wt_: EHA105 wild-type DNA control; P_E_: pDONR1K18-EHAmetA plasmid; W: water; Lanes 9 to 16 LBA4404 background colony screening; L_wt_: LBA4404 wild-type DNA control; P_L_: pDONR1K18-LBAmetA plasmid; L: 100-bp DNA ladder. *: mutant allele at *met*A locus (381 and 455-bp for EHA105_met_ and LBA4404_met_, respectively). ◀: wild-type allele (1,271 and 1,349-bp for EHA105 and LBA4404, respectively). (C) Methionine dependent growth of auxotrophic strains after 24-hour incubation at 28 °C in minimal medium (M9) or M9 supplemented with DO.

The vector, pCAMBIA-1305.2 (Jefferson et al., 2002), was electroporated at 1 ng µL^-1^ into the *Agrobacterium* strains. Bacteria were allowed to recover for 2-4 hours before plating either on solid YM medium (for parental/WT strains) or solid YM plus 0.77 g L^-1^ DO -Ura supplement for auxotrophic strains, both supplemented with 50 mg L^-1^ each of rifampicin and kanamycin for EHA105 and 50 mg L^-1^ kanamycin with 250 mg L^-1^ streptomycin for LBA4404. Colonies were PCR-screened to confirm presence of the plasmid and the mutation. Standard PCR was performed using the Apex 2X Taq RED Master Mix (Genesee Scientific, San Diego, CA, USA), with 1 µL of bacterial culture at OD_600_ 1.0 and primers at 0.25 µM. PCR conditions were initial denaturation at 94 °C for 4 min, 35 cycles of (1) denaturation at 94 °C for 30 sec (2) annealing temperature for 30 sec, and (3) extension at 72 °C for 1 min per kb, and final extension at 72 °C for 7 min. Primers were designed using Geneious^®^ 11.1.5 (https://www.geneious.com) and are in Table S1.

### Switchgrass *in vitro* plantlets and culture of explants

Switchgrass (*Panicum virgatum* L.) Performer 7 (Ondzighi-Assoume et al., 2019) *in vitro* plants were propagated aseptically in Nexclear^®^ cups (0.6 L) with dome lids (Fabri-Kal^®^, Kalamazoo, MI, USA). The medium half-strength MS medium (Murashige and Skoog, 1962) supplemented with B5 vitamins (Gamborg et al., 1968) and 1.5% w/v sucrose (½ MS-B5) at 26 °C, with cool-white fluorescent lighting (66-95 μE m^-2^ s^-1^) and a 23-h light: 1-h dark photoperiod. Every 4-6 weeks, newly emerging tillers from plantlets were subcultured for propagation.

The culture procedure was modified from King et al. (2014). Four-week-old *in vitro* plants, about 10-cm tall, were used as starting material for callus induction. Leaves were removed and intact stems were placed in a 20 × 145-mm Petri dish, and the 0.5-cm-long segment was removed from the basal culm. The basal culm is the colorless zone at the base above the roots, just above the roots. The basal culms were halved longitudinally and placed cut-side down on MS medium with MS vitamins supplemented with 5 mg L^-1^ 2,4-dichlorophenoxyacetic acid (2,4-D), 0.15 mg L^-1^ 6-benzylaminopurine (BAP) and 30 g L^-1^ sucrose (MSD5B0.15) for callus induction. Explants were arranged in a 5 × 5-grid pattern in 15 × 100 mm Petri dishes sealed with 3M™ Micropore™ tape (3M Health Care, Saint Paul, MN, US). Tissue was incubated at 28 °C in the dark for three weeks.

Proliferating calli were transferred to switchgrass callus maintenance medium consisting of MS medium supplemented with MS vitamins, 5 mg L^-1^ 2,4-D, 1 mg L^-1^ BAP and 30 g L^-1^ maltose (MSD5B1) in a 5 × 5-grid pattern in 15 × 100 mm Petri dishes and incubated for an additional three weeks as described above. At the end of this period, only type II embryogenic callus (Burris et al., 2009) was selected and bulked with subcultures at three-week intervals in MSD5B1.

Embryogenic calli (∼5 mm diameter pieces) were transferred to MS medium supplemented with 1 mg L^-1^ BAP and B5 vitamins (Alexandrova et al., 1996) but with 30 g L^-1^ sucrose and pH 5.8 for shoot regeneration (RMS-B1, King et al., 2014). Calli were arranged in a 5 × 5-grid pattern in 20 × 100 mm Petri dishes sealed with 3M™ Micropore™ tape. Calli were incubated at 26 °C, with cool-white fluorescent lighting (66-95 μE m^-2^ s^-1^) and a 23-h light: 1-h dark photoperiod for two weeks. After that, calli were transferred onto fresh RMS-B1 and incubated for an additional 2 weeks. Emerging shoots longer than 0.5 cm were rooted Nexclear^®^ cups as described for the explant source. Root and shoot elongation were visible after 4-6 weeks (Figure 2).

**Figure 2.**
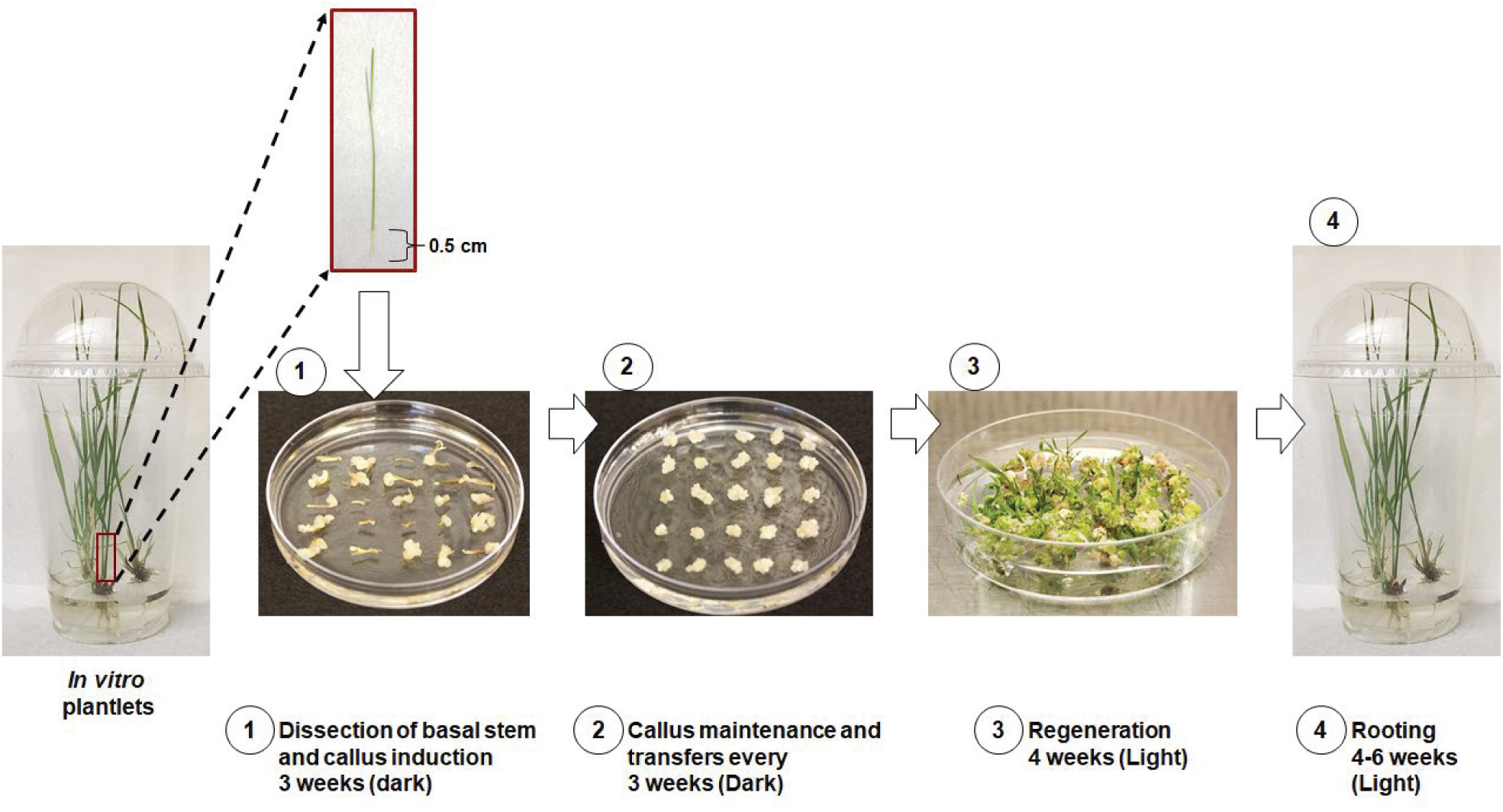
Embryogenic callus induction from basal culms of *in vitro* switchgrass plantlets.

All the media used in switchgrass tissue culture, except RMS-B1, were solidified with 2.5 g L^-1^ Gelzan™ (Bioworld, Dublin, OH, USA). RMS-B1 was solidified with 2.5 g L^-1^ Phytagel™ (Sigma-Aldrich, St. Louis, MO, USA). All medium pH was adjusted to pH 5.8 with HCl or NaOH before autoclaving (Table S3).

### Rice plant material and culture of explants

Rice (*Oryza sativa* subspecies japonica cv. Taipei 309, TP309) seeds were surface-sterilized as described by (Mann et al., 2011), with 70% ethanol for 2 min with manual agitation. Kernels were then submerged in a 60% Clorox^®^ solution (v/v) supplemented with two drops of Tween-20 per 100 mL and swirled for 30 min at 100 rpm on a shaker. Kernels were washed three times with sterile deionized water for 2-min. Once sterilized, kernels were dried on autoclaved Whatman^®^ Grade 1 qualitative filter paper (Cytiva, Marlborough, MA, USA). Dried kernels were organized in a 5 × 5-grid on modified NB (mNB) medium (Chen et al., 1998) supplemented with 2 mg L^-1^ 2,4-D for initiation and maintenance of embryogenic callus. Tissue was incubated at 28 °C in the dark, and compact embryogenic callus pieces were selected and transferred to fresh mNB medium every 3 weeks. For shoot regeneration, embryogenic calli were transferred to N6 medium (Chu et al., 1975) supplemented with N6 vitamins, 3 mg L^-1^ BAP and 0.5 mg L^-1^ 1-naphthaleneacetic acid (NAA) (named RGH6) and placed in the dark at 28 °C for 7 days. After one week, cultures were transferred to the light under a 23 h-light/1 h-dark regime provided by cool-white fluorescent light (66-95 µE m^-2^ s^-1^) and incubated at 26 °C for up to six weeks. Emerging shoots (≤1 cm tall) were rooted in ½ MS supplemented with B5 vitamins and hygromycin at 50 mg L^-1^ (½ MS-B5).

All the media used in rice tissue culture, except RGHG, were solidified with Gelzan™. Shoot regeneration medium (RGHG) was solidified with 2.5 g L^-1^ Phytagel™. Gelzan was used at 2.5% in callus induction and maintenance medium (mNB) and 3% in rooting medium (½ MS-B5). All medium pH was adjusted to pH 5.8 with HCl before autoclaving (Table S3).

### Evaluation of transient transformation efficiency

Three days prior to transformation, embryogenic callus 5 mm in size were transferred to either fresh MSD5B1 or mNB callus maintenance medium for each plant species as previously described.

Bacterial strains were streaked on YM solid medium two days before tissue transformation. *Agrobacterium* was grown on solid YM supplemented with the proper antibiotics for bacteria and plasmid selection. Rifampicin and kanamycin at 50 mg L^-1^ were used for selection in EHA105 and its auxotrophic strain, and 250 mg L^-1^ streptomycin and 50 mg L^-1^ kanamycin were used for LBA4404 and its derived auxotrophic strain. For auxotrophic strains, YM was supplemented with DO supplement -Ura. Bacterial plates were incubated at 28 °C in the dark. Two-day-old *Agrobacterium* colonies were picked and resuspended in liquid mNB supplemented with 100 µM acetosyringone (Sigma-Aldrich, St. Louis, MO, USA). Each bacterial strain was adjusted to a final concentration of OD_600_ ∼0.5 in a 50-mL sterile tube and incubated at 21 °C for 90 min with continuous agitation at 220 rpm.

Embryogenic calli were placed in 50-mL sterile tubes and mixed with 2.5 – 3 mL of the *Agrobacterium* liquid culture and allowed to sit at room temperature for 15 min. Callus pieces were dried for long enough to remove the excess liquid on three disks of glass fiber filter (VWR, Radnor, PA, USA) and placed into a 15 × 100 mm Petri dish. Callus pieces were placed in a 5 × 5-grid pattern on mNB supplemented with 100 µM acetosyringone in 15 × 100 mm Petri dishes sealed with 3M™ Micropore™ tape. Petri dishes were covered with aluminum foil before incubation at 21°C for either 3, 4, or 5 days. Co-cultivated calli were initially rinsed eight times with autoclaved deionized water at 1-minute intervals with gentle manual agitation. Tissue was washed twice in sterile deionized water with meropenem (TCI chemicals, Tokyo, Japan) at 50 mg L^-1^ for EHA105 and derivative strains or 10 mg L^-1^ for LBA4404 and derivative strains. Callus pieces were blotted dry as before. Callus pieces were transferred to mNB medium supplemented with meropenem as previously described in 15 × 100 mm Petri dishes sealed with 3M™ Micropore™ in a 5 × 5-grid pattern and incubated at 28 °C in darkness for 8 days. GUS assays were conducted in callus pieces according to Jefferson (1987). Tissue was then incubated at 37 °C for 24 hours and washed three times with 70% ethanol at 24-hour intervals to remove the GUS staining buffer. Pieces of GUS-stained callus were visualized on an Olympus SZX12 Stereo Microscope equipped with an Olympus DP27 Camera (Olympus, Center Valley, PA, USA). GUS-positive areas were quantified with ImageJ version 1.53a (Schneider et al., 2012).

### Switchgrass and rice stable transformation

Stable transformation was achieved by following the same steps as previously described for transient expression. Following the 8-day resting period, calli were transferred to MSD5B1 supplemented with 200 mg L^-1^ hygromycin B and meropenem as described previously. Each callus was broken up into smaller 1-2 mm diameter pieces as described by Chen et al. (1998) to ensure independent events were recovered. Incubation was at 28 °C in dark conditions for 8 weeks with one subculture at 4 weeks.

### Molecular analysis of regenerated events

Genomic DNA (gDNA) from switchgrass leaves was extracted with a modified chloroform CTAB procedure (Stewart and Via, 1993). Frozen leaf samples (∼0.1 g) were ground with 900 µL CTAB buffer with two 4.8 mm steel beads (Med Supply Partners, Atlanta, GA, USA) in a GenoGrinder 2010 (Spex SamplePrep, Metuchen, NJ, USA) for 2 min at 1,600 strokes min^-1^. Samples were incubated for 30 min at 65 °C. 800 µL chloroform:isoamyl alcohol (24:1) were added and the water-soluble fraction was recovered after centrifugation for 10 min at 16,000 x g. DNA was precipitated with 100% isopropanol and centrifugation as described before. Finally, DNA was washed twice with 70% ethanol and resuspended in 50 µL of TE buffer pH 8.0 and incubated at 42 °C for 10 min to dissolve the pellet. DNA was quantified on a Synergy™ 2 microplate reader (Biotek Instruments Inc., Winooski, VT, USA) and diluted to 20 ng μL^-1^ with Type I water.\ Standard PCR was performed as described previously. Primers to amplify a 596 bp region within *hph* were used (Table S1, Figure S1). PCR was also used to check for the presence of residual *Agrobacterium* in plant gDNA using primer combination EHA105_metA_seqF/EHA105_metA_seqR and LBA4404_metA_seqF1/LBA4404_metA_seqR1 (Table S1). Non-transgenic template DNA, pCAMBIA-1305.2 plasmid DNA, water, and *Agrobacterium* strains from this study were used as controls. PCR products were visualized on a 0.5xTBE 1% agarose gel supplemented with 0.025 μL mL^-1^ of MIDORI^Green^ Xtra stain (NIPPONgenetics, Dueren, Germany).

Digital droplet PCR (ddPCR) was used for transgene copy number detection. No-template control samples were also included in all the runs to surveil for the appearance of false positive droplets. Briefly, 2 μg gDNA were digested with 40 U of *Xho* I restriction enzyme (New England Biolabs Inc., Ipswich, MA, USA) in a final 50 μL restriction dilution and incubated at 37 °C for 3 h. After purification of digested gDNA with a DNA Clean & Concentrator^*TM*^ (Zymo Research Corp., Irvine, CA, USA), a 22-μL ddPCR reaction mix was prepared with 5.5 ng of purified digested gDNA, 900 nM of each primer (for both *GUSPlus*^*TM*^ and reference gene), 250 nM of each probe and 11 μL of 2x ddPCR Supermix for Probes (No dUTP) (BioRad, Hercules, CA, USA). Droplets were generated according to the manufacturer’s instructions in a QX200™ AutoDG Droplet Generator (BioRad, Hercules, CA, USA) using 20 μL of ddPCR reaction mix and 20 μL of droplet generation oil for probes (BioRad, Hercules, CA, USA). Generated droplets (40 μL) were loaded onto a 96-well PCR plate (Eppendorf, Enfield, CT, USA) and PCR was conducted according to the following conditions: initial denaturation at 95 °C for 10 min, 40 cycles of (1) denaturation at 94 °C for 30 sec and (2) annealing/extension at 60 °C for 1 min with a temperature ramp rate of 2 °C s^-1^ between all temperatures according to manufacturer guidelines, and final step at 98 °C for 10 min. Primers and probe were designed using PrimerQuest design tool from IDT Technologies (https://www.idtdna.com), and those are described in Table S1. After thermocycling, PCR was analyzed in a QX200 droplet reader (BioRad, Hercules, CA, USA).

Calculation of the absolute concentration of PCR template using fluorescence readings was performed in QuantaSoft software™ version 1.7.4.0917 (BioRad, Hercules, CA, USA), where the population of negative and positive droplets determines the initial template concentration according to the Poisson probability distribution. The ratio between the number of copies per nanogram of digested genomic DNA of the *GusPlus*^*TM*^ transgene and concentration of reference gene was used to calculate copy number. The purine nucleoside phosphorylase gene (PNP, Pavir.3KG080700) was selected as a reference gene for switchgrass as found a single copy gene and specific primers were designed to amplify 115 bp. The *GusPlus*^*TM*^ and the reference gene probes were labeled with the fluorescent hexachloro-fluorescein (HEX™) and carboxy-fluoroscein (FAM™), respectively. Droplets were generated and PCR performed by University of Florida ICBR Gene Expression and Genotyping Core Facility (RRID:SCR_019145).

### Experimental design & statistical analyses

All transformation experiments (transient and stable) used a randomized complete block design with three (transient transformation) or two replicate (stable transformation) plates per treatment. Each treatment consisted of an *Agrobacterium* to be evaluated and three different co-cultivation times consisting of 3, 4 or 5 days. Each experimental unit consisted of a 15 × 100-mm Petri dish containing 25 embryogenic calli.

For transient expression, the percentages of GUS-stained surfaces were transformed (square root) and subjected to Shapiro-Wilk Test to check for homogeneity of variance using GraphPad Prism version 9.2.0 for Windows (GraphPad Software, San Diego, CA, USA). Data were normally distributed, so they were subjected to a two-way ANOVA to analyze significant differences between treatments within an experiment (p ≤ 0.05) and a Tukey test to determine which groups are different (*post hoc*).

Stable transformation was tabulated as the number of hygromycin B-resistant calli observed after eight weeks of selection. Data were tested for homogeneity of variance. Data were normally distributed (p > 0.05), so they were subjected to a two-way ANOVA without interaction (only main effects) to detect significant treatment effect (p ≤ 0.05 and a Tukey HSD test was done to look for significant differences within treatments (p ≤ 0.05).

Digital droplet PCR statistical analysis and plots were performed with the R open-source software (version 4.1.1; R Core Team 2021).

## Results and discussion

### Generation of auxotrophic strains

Standard cloning techniques were used (see Methodology section) to delete the homoserine O-succinyltransferase (*metA*) gene, in *Agrobacterium tumefaciens* EHA105 and LBA4404. After transformation with the deletion construct, followed by selection and counterselection, eight colonies for each strain were screened using PCR to verify deletion of the *metA* gene in all copies of the *A. tumefaciens* chromosome (Figure 2). For EHA105, two colonies (#2 and #5; Figure 2B – lane 2 and 5) had only the amplicon for the gene deletion with no amplicon for the wild-type chromosome. For LBA4404, we identified a single colony (#5; Figure 2B lane 13) that was a segregated deletion strain. Freezer stocks of each colony were made by growing the colonies overnight in YM medium and then diluting them with 75% (v/v) glycerol for a final concentration of 15% glycerol. Each colony was then grown in M9 minimal medium and M9 minimal medium supplemented with methionine (the methionine was provided by adding an amino acid dropout mix, DO supplement –Ura). Strains auxotrophic for methionine are unable to grow in M9 medium without supplemental methionine. The EHA105 Δ*metA* colony #2 did not consistently show auxotrophic growth or correct PCR amplicons. In contrast, EHA105 Δ*metA* colony #5 and LBA4404 Δ*metA* colony #5 both consistently exhibited auxotrophic growth and were therefore used for further study. They are henceforward referred to as EHA105_met_ and LBA4404_met_, respectively.

### The use of in vitro plants for explants

When using *in vitro* plants, the bottom 0.5-cm segment of the basal culm provides the best response for callus initiation (Denchev and Conger, 1994; Denchev and Conger, 1995). As contamination is a major problem when explants are taken from greenhouse-grown plants, *in vitro*-produced aseptic plants were used (Leifert et al., 1994). Importantly, the use of *in vitro* clonally propagated plantlets as the explant donor allows for maintenance of genotype fidelity, which would be lost if seed-derived explants were used. Finally, the *in vitro* plants are much smaller in size than greenhouse-grown plants, requiring less greenhouse space for plant maintenance.

### GUS staining after co-cultivation

Switchgrass and rice calli were inoculated for 3, 4, or 5 days with EHA105, EHA105_*met*_, LBA4404, or LBA4404_*met*._, and then screened for GUS expression (Figure 3; Figure 4). Based on a two-way ANOVA, strain and co-cultivation time affected results (P < 0.01) as did their interaction (P < 0.05) in both crops. In switchgrass, EHA105 and its auxotroph with a minimum of four days of co-cultivation gave better response than LBA4404 and its auxotroph (Figure 4).

**Figure 3.**
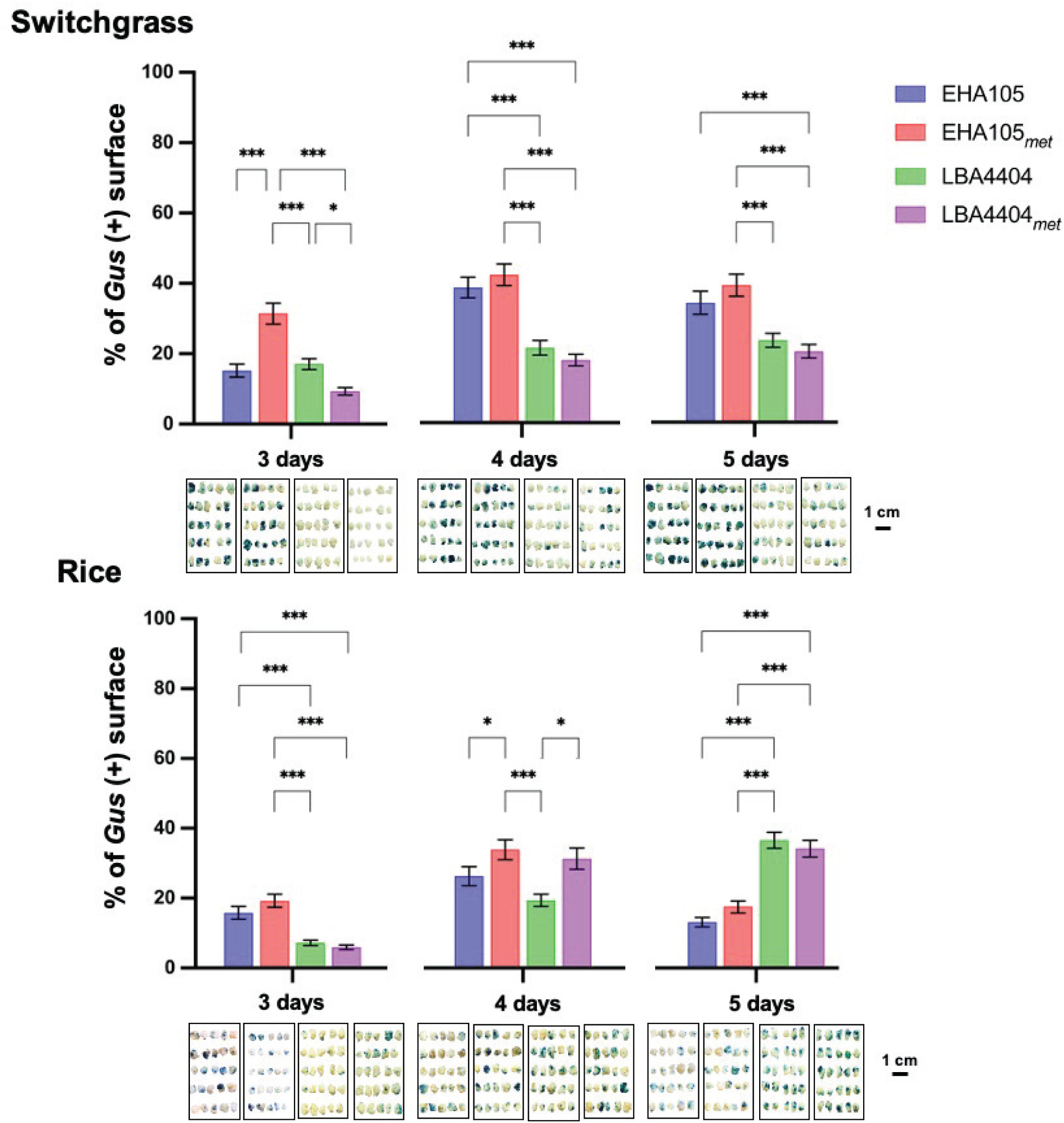
GUS staining in Switchgrass Performer 7 (top) rice TP309 (bottom) callus including all *Agrobacterium* strains used in this study at three, four, and five days of co-cultivation. Photos from one rep are shown. Plot and error bars represent the mean value ± SEM of percent GUS staining from three biological replicates (n=25). Only significant pairwise comparisons are shown. Significance levels P-value according to Tukey test are shown as <0.05 (*), <0.01 (**), or <0.001 (***).

**Figure 4.**
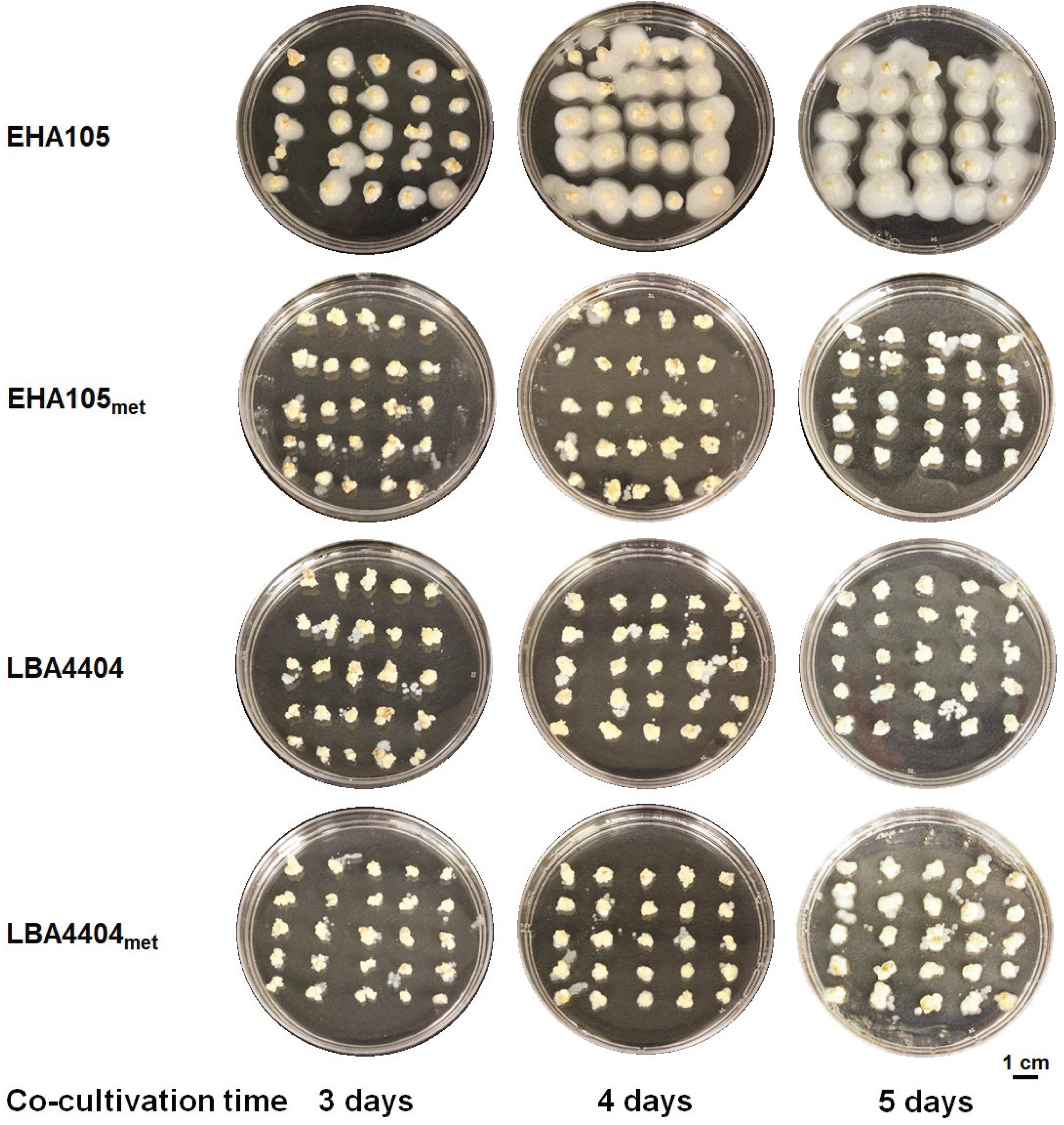
*Agrobacterium* growth in co-cultivation with switchgrass tissue. Co-cultivation was set as three, four and five days.

**Figure 5.**
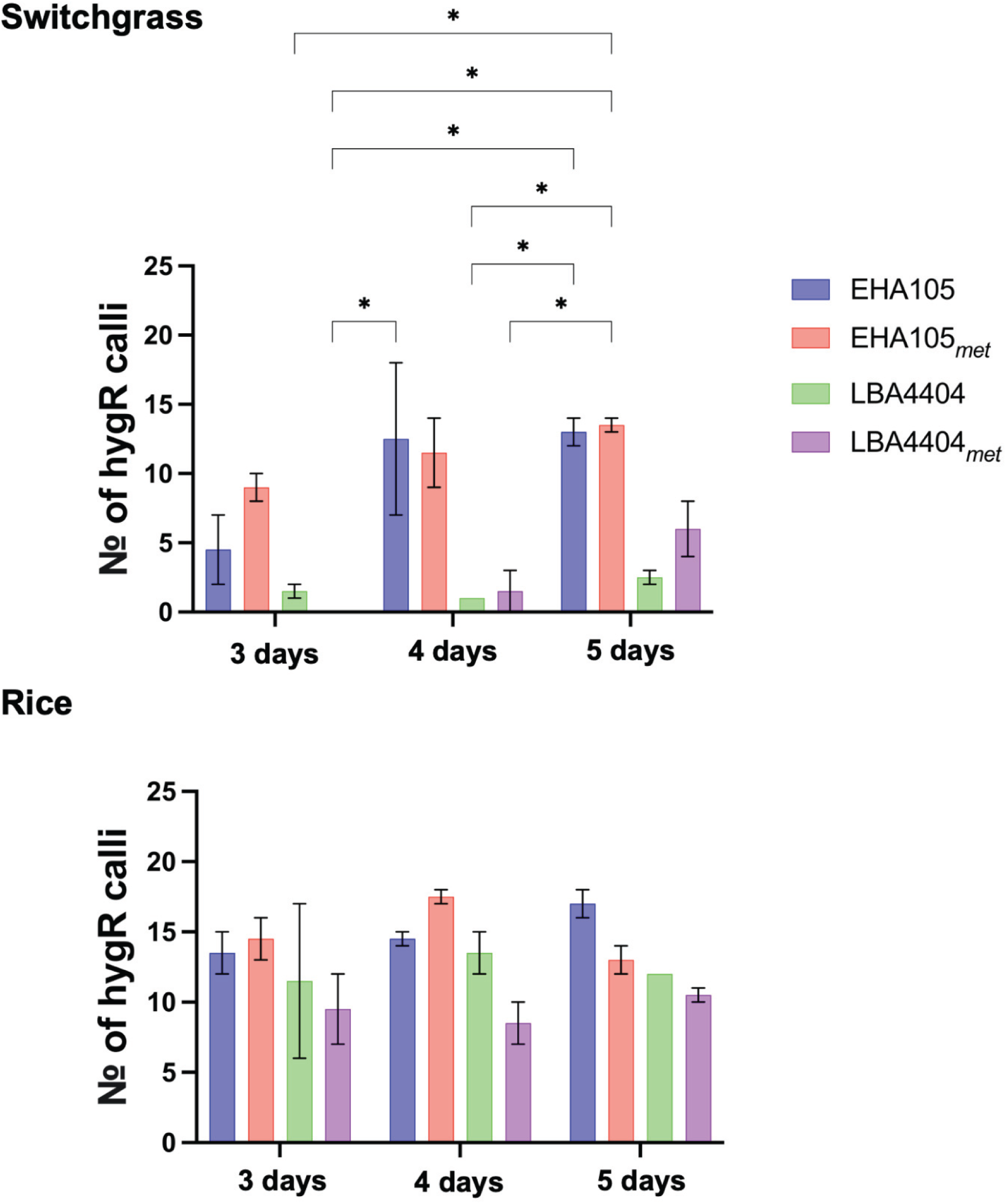
Number of hygromycin resistant callus after transformation of switchgrass and rice with EHA105 and EHA105_met_, and LBA4404 and LBA4404_met_. Bars represent the mean ± SEM of hygromycin B resistant calli from two biological replicates after eight weeks of selection. Significance levels P-value according to Tukey’s test are shown as > 0.05 (ns), ≤ 0.05 (*), ≤ 0.01 (**), or ≤ 0.001 (***).

For rice, the two auxotrophic strains gave the best results at four days of co-cultivation (Figure 4). EHA105_*met*_ with three days of co-cultivation was significantly better than LBA4404 and LBA4404_*met*_ (Figure 4). However, by 5 days of co-cultivation both LBA4404 and LBA4404_*met*_ showed the best response (Figure 4).

The most notable difference is the tendency of EHA105 to overgrow the tissues during co-cultivation. Overgrowth is not as much of an issue with its auxotroph or the LBA4404 strains. The high transformation capacity of EHA105 for switchgrass makes this strain one of the most used for transgenic switchgrass production; however, antibiotics are required at high concentrations to control bacterial overgrowth on the plant tissue (Lin et al., 2017). In addition, overgrowth of *Agrobacterium* and high concentrations of antibiotics have been known to either reduce transformation efficiency or affect plant cell differentiation (Liu et al., 2016).

### Efficiency of transformation

In general, the amount of GUS staining 8 days after co-cultivation was indicative of the amount of hygromycin-resistant callus that was obtained later for switchgrass but not for rice. The actual growth of transgenic callus is a better indicator of transformation than GUS staining. For switchgrass, the highest rates of hygromycin-resistant calli were obtained with the EHA105 strains after 4 and 5 days of co-cultivation. As evidenced by the smaller error bars, transformation rates were more uniform with 5 days of co-cultivation. This observation is consistent with that of a previous study in which EHA105 induced high levels of GUS expression in switchgrass embryogenic callus and seedlings when compared to strains LBA4404 and AGL1 (Xi et al., 2009).

The length of the co-cultivation time has been directly correlated with transformation efficiencies in ‘Alamo’ switchgrass, such that a 7-day co-cultivation yielded 3.3-fold higher transformation than did a co-cultivation of three days (Ogawa et al., 2014). The results here suggest a similar trend, and tissues subjected to longer co-cultivation periods were more uniformly transformed. Thus, it may be feasible to get even higher levels of transformation with prolonged days in co-cultivation and without tissue overgrowth (Cervera et al., 1998; Kondo et al., 2000).

The difference between strains is less apparent for rice. Even 3 days of co-cultivation resulted in reasonable transformation rates. With additional co-cultivation time, the EHA strains have a small but significant advantage over the LBA4404 strains.

### Quality of transformation events

Besides transformation efficiency, the amount of T-DNA integrated into the transgenic events is important. Ideally, transgenics should have just 1 copy of the T-DNA. As the effect of auxotrophy on amount of T-DNA delivered to transgenics has not been published, we set to evaluate the quality of the recovered switchgrass events. High quality transformation events are those containing a single copy insertion of the complete, intact T-DNA and have no binary vector backbone (Harwood, 2012).

A total of 107 plants (95 EHA105 strains and 12 LBA4404 background) were regenerated and evaluated by droplet digital PCR (ddPCR). We used GusPlus™ as the target sequence to determine transgene presence and the purine nucleoside phosphorylase gene (PNP) as the single-copy gene to use as endogenous reference control gene and assessed transgene copy number (Figure 6). On average EHA105 and EHA105_met_ delivered similar number of transgene copies (2.37 and 2.33 copies, respectively). The number of T-DNA sequences delivered by EHA105 and EHA105_met_ ranged from 1 to 7 copies, with one case in which EHA105_met_ delivered up to 12 copies. No significant differences in quality were identified across days of co-cultivation.

**Figure 6.**
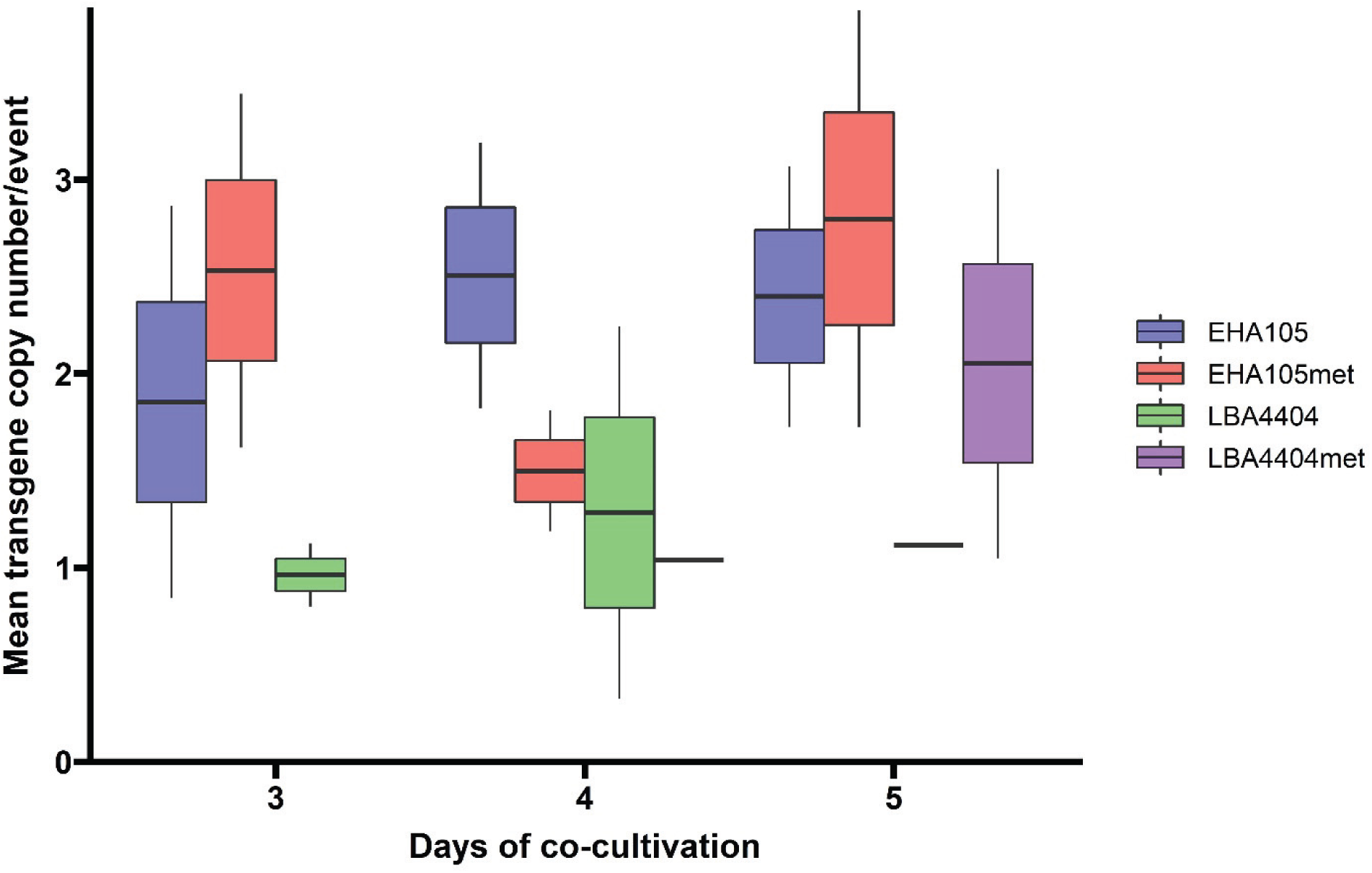
Evaluation of transgene copy number among switchgrass transgenic T_0_ lines transformed with pCAMBIA-1305.2. Box plots depict the mean ± SEM of GusPlus™ copies inserted by the different *Agrobacterium* strains, with whiskers showing the 95%-confidence interval. Each *Agrobacterium* strain is colored as indicated in the key. The low mean value at 4-days of co-cultivation for EHA105met: was because 9 of the 15 events analyzed (60%) presented one single T-DNA insertion.

In contrast, plants transformed with LBA4404 strains display a lower number of insertions: LBA4404 delivered between 1 and 2 transgene copies whereas LBA4404_met_ activity ranged from 0 to 5 copies at 3 and 5-days of co-cultivation, respectively. On average, LBA4404 inserted 1.12 copies and LBA4404_met_ inserted 1.91 copies of GusPlus™.

Auxotrophy has no effect on the number of T-DNA copies delivered by any of the *Agrobacterium* strains evaluated. These results support those of Zhi et al. (2015), who also found that the use of LBA4404 resulted in fewer but higher quality transgenic events than the use of AGL0, a sister strain to EHA105.

These results are consistent with those of the recent literature that show that knocking out genes to cause auxotrophy does not affect transformation capacity. Nevertheless, auxotrophy in *Agrobacterium* historically has been associated with loss of virulence, and this original negative perception may have discouraged researchers from using auxotrophic strains. Garber et al. (1955) summarized the variable virulence of three *Agrobacterium* auxotrophic strains and reported a very low or lost virulence for the tryptophan mutant. Left unsaid was that since UV mutagenesis produces random mutations, other genes that affect virulence could have been inadvertently mutated, accounting for the low virulence of some auxotrophs (Garber et al., 1955).

Lippincott et al. (1965) also evaluated the effect of auxotrophy on tumor-inducing capacity in primary bean leaves. Six auxotrophic strains were developed either by UV light irradiation or heat treatment at 42-45 ⁰C. Infectivity of the auxotrophs ranged from 1-30% of that of the parent strain (set as 100% infectivity). However, the results are based on a low number of samples with only one plant infected per auxotroph, and with high variability between the same treatment in different experiments. (Lippincott et al., 1965). A year later, the same authors showed that infectivity of the auxotrophic strains could be increased by supplying the missing nutrient to the pinto bean leaves prior to infection (Lippincott and Lippincott, 1966).

Overall, the use of auxotrophic strains of *A. tumefaciens* minimizes the use of antibiotics to prevent overgrowth of the plant tissue (Dirks and Peeters, 2001; Ranch et al., 2010). To wit, the recent use of morphogenic genes for monocot transformation, mainly maize, was facilitated using an LBA4404 thymidine auxotroph (Lowe et al., 2018; Lowe et al., 2016) with the advantage of a reduced/controlled bacterial growth in tissue culture (Ranch et al., 2010). No doubt the use of auxotrophic strains will facilitate additional applications in the future once auxotrophic strains are available to researchers in public institutions.

## Acknowledgements

Funding was provided by The Center for Bioenergy Innovation, a U.S. Department of Energy Research Center supported by the Office of Biological and Environmental Research in the DOE Office of Science.

## Supplementary Information

**Table S1:**
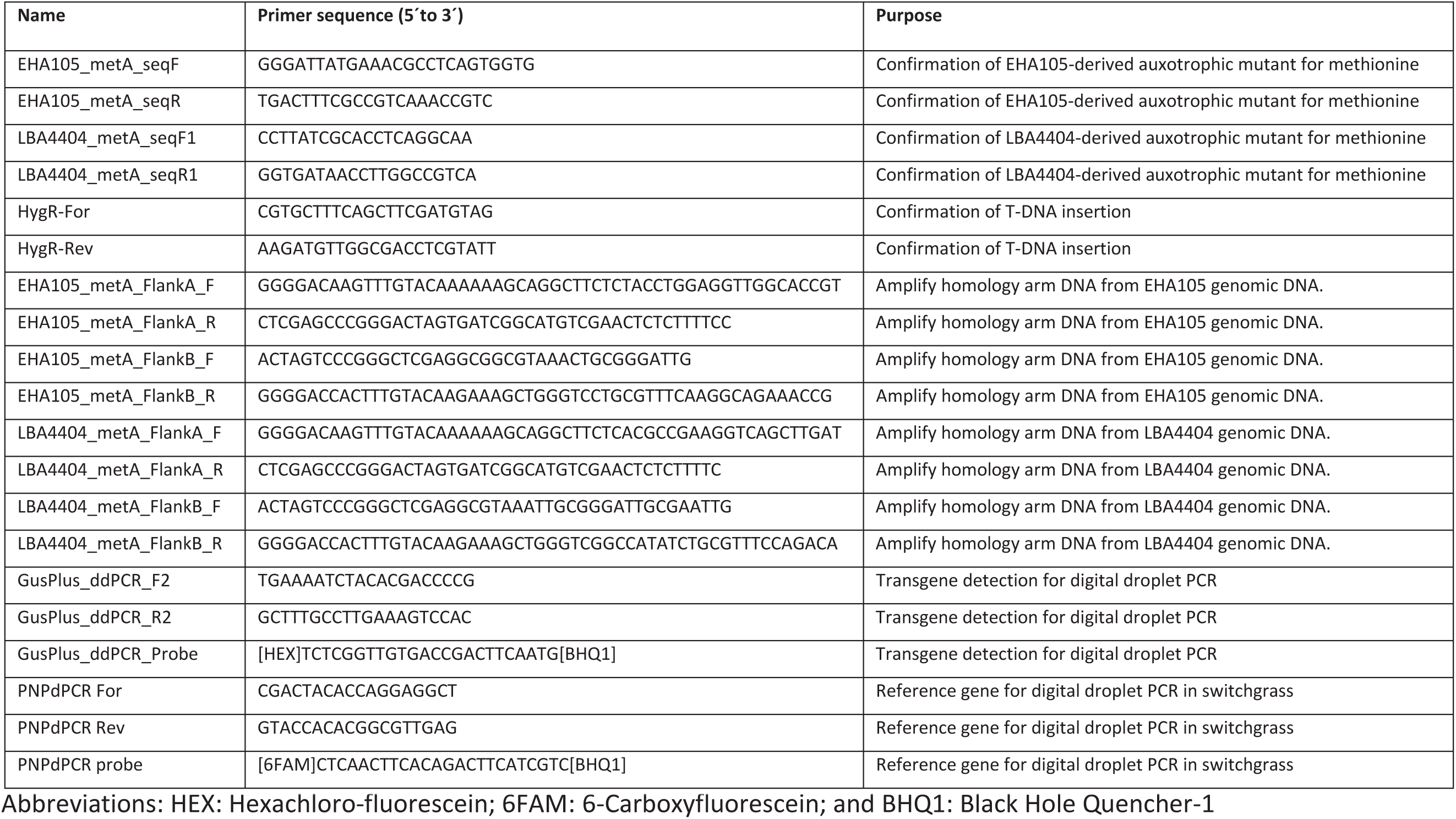
Oligonucleotides used for assembling suicide plasmids and to identify and screen for auxotrophic *Agrobacterium* strains and pCAMBIA-1305.2 plasmid.

**Table S2:** Regions containing the gene necessary for metA biosynthesis in EHA105 and LBA4404.

| Strain | Deleted gene | Gene product | Locus tag | GenBank | Deficiency |
| --- | --- | --- | --- | --- | --- |
| EHA105 | metA | Homoserine O-succinyltransferase | Atu2718 | NC_003062 (2703611...2704537) | Methionine |
| LBA4404 | metA | Homoserine O-succinyltransferase | Ach5_RS13190 | NZ_CP011246 (2696107...2697033) | Methionine |

**Table S3:**
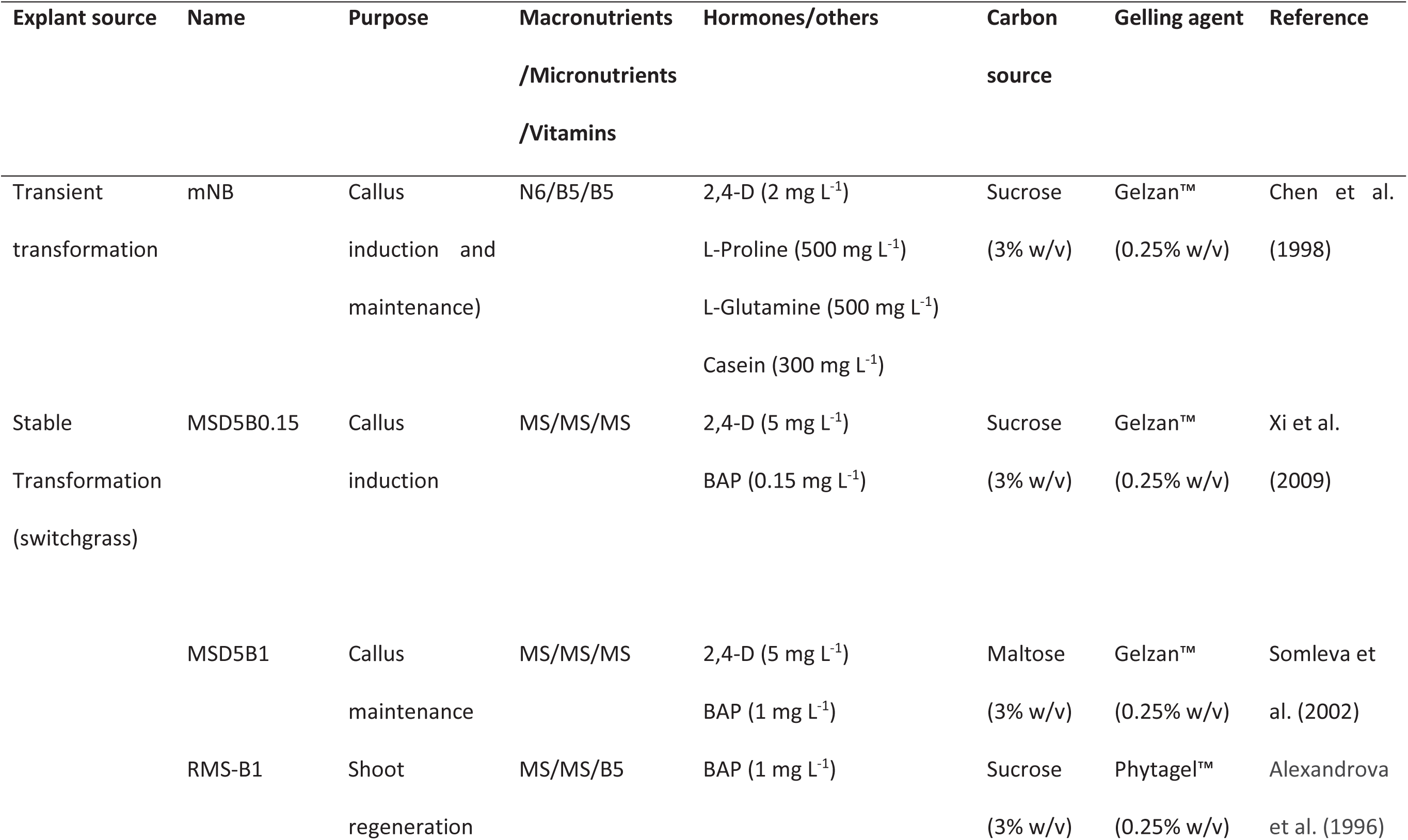
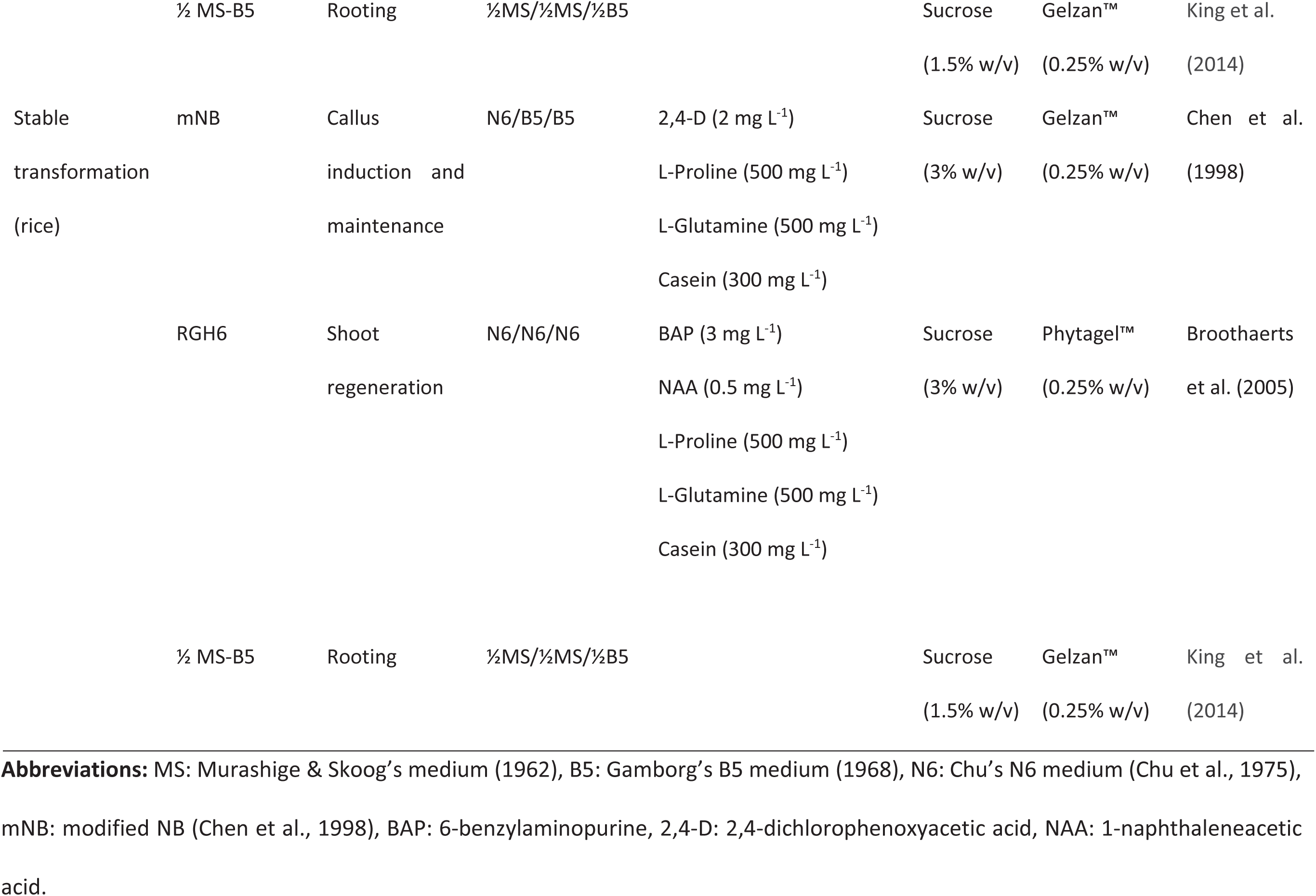
Media used in tissue culture and transformation of Performer 7 switchgrass and TP309 rice.

**Figure S1.**
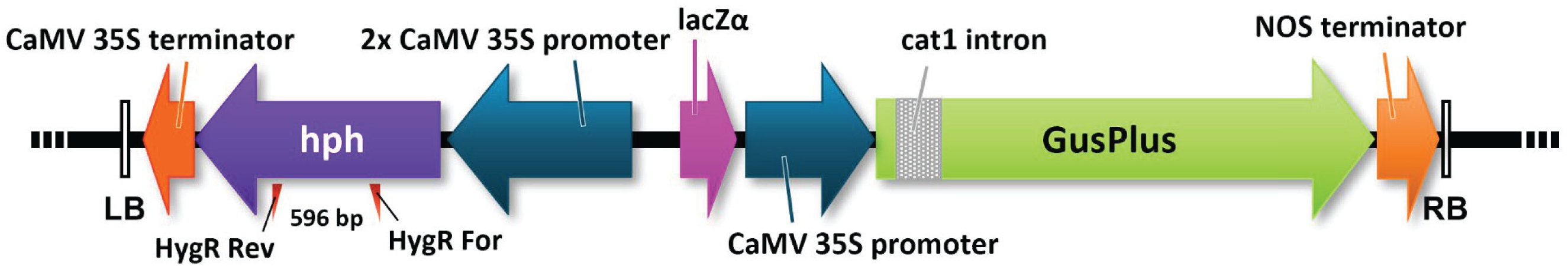
Schematic representation of T-DNA region of binary vector pCAMBIA-1305.2. Red triangles indicate the relative position of the PCR primers used to confirm T-DNA insertion. The expected PCR product size is indicated in space between the two primers

## REFERENCES

Alexandrova, K.S., Denchev, P.D. and Conger, B.V. (1996) In vitro development of inflorescences from switchgrass nodal segments. Crop Science 36: 175–178.

Aliu, E., Azanu, M.K., Wang, K. and Lee, K. (2020) Generation of thymidine auxotrophic Agrobacterium tumefaciens strains for plant transformation. bioRxiv. doi: 10.1101/2020.08.21.261941.

Anand, A., Bass, S.H., Wu, E., Wang, N., McBride, K.E., Annaluru, N., Miller, M., Hua, M. and Jones, T.J. (2018) An improved ternary vector system for Agrobacterium-mediated rapid maize transformation. Plant Molecular Biology 97: 187–200.

Broothaerts, W., Mitchell, H.J., Weir, B., Kaines, S., Smith, L.M.A., Yang, W., Mayer, J.E., Roa-Rodríguez, C. and Jefferson, R.A. (2005) Gene transfer to plants by diverse species of bacteria. Nature 433: 629–633.

Burris, J.N., Mann, D.G.J., Joyce, B.L. and Stewart, C.N. (2009) An improved tissue culture system for embryogenic callus production and plant regeneration in switchgrass (Panicum virgatum L.). BioEnergy Research 2: 267–274.

Cervera, M., Pina, J.A., Juárez, J., Navarro, L. and Peña, L. (1998) Agrobacterium-mediated transformation of citrange: factors affecting transformation and regeneration. Plant Cell Reports 18: 271–278.

Chen, L., Zhang, S., Beachy, R.N. and Fauquet, C.M. (1998) A protocol for consistent, large-scale production of fertile transgenic rice plants. Plant Cell Reports 18: 25–31.

Chu, C.-C., Wang, C.-C., Sun, C.-S., Hsu, C., Yin, K.-C. and Bi, F.-Y. (1975) Establishment of an efficient medium for anther culture of rice through comparative experiments on the nitrogen sources. Scientia Sinica 18: 659–668.

Collens, J.I., Lee, D.R., Seeman, A.M. and Curtis, W.R. (2004) Development of auxotrophic Agrobacterium tumefaciens for gene transfer in plant tissue culture. Biotechnology Progress 20: 890–896.

Collens, J.I., Mason, H.S. and Curtis, W.R. (2007) Agrobacterium-mediated viral vector-amplified transient gene expression in Nicotiana glutinosa plant tissue culture. Biotechnology Progress 23: 570–576.

Denchev, P.D. and Conger, B.V. (1994) Plant regeneraton from callus cultures of switchgrass. Crop Science 34: 1623–1627.

Denchev, P.D. and Conger, B.V. (1995) In vitro culture of switchgrass: influence of 2, 4-D and picloram in combination with benzyladenine on callus initiation and regeneration. Plant Cell Tissue and Organ Culture 40: 43–48.

Dirks, R. and Peeters, R. (2001) Agrobacterium-mediated transformation of plants. United States Patent number 6, 323, 396.

Gamborg, O.L., Miller, R.A. and Ojima, K. (1968) Nutrient requirements of suspension cultures of soybean root cells. Experimental Cell Research 50: 151–158.

Garber, E.D., Goldman, M. and Hirshhorn, S.G. (1955) Auxotrophy and virulence in Agrobacterium tumefaciens. In: Proceedings of the 55th General Meeting p. 122. New York: The Society of American Bacteriologists.

Harwood, W.A. (2012) Advances and remaining challenges in the transformation of barley and wheat. Journal of Experimental Botany 63: 1791–1798.

Hoekema, A., Hirsch, P.R., Hooykaas, P.J.J. and Schilperoort, R.A. (1983) A binary plant vector strategy based on separation of vir-and T-region of the Agrobacterium tumefaciens Ti-plasmid. Nature 303: 179–180.

Hoerster, G., Wang, N., Ryan, L., Wu, E., Anand, A., McBride, K., Lowe, K., Jones, T. and Gordon-Kamm, B. (2020) Use of non-integrating Zm-Wus2 vectors to enhance maize transformation. In Vitro Cellular & Developmental Biology-Plant: 1–15.

Hood, E.E., Gelvin, S.B., Melchers, L.S. and Hoekema, A. (1993) New Agrobacterium Helper Plasmids for Gene-Transfer to Plants. Transgenic Research 2: 208–218.

Jefferson, R.A. (1987) Assaying chimeric genes in plants: the GUS gene fusion system. Plant Molecular Biology Reporter 5: 387–405.

Jefferson, R.A., Harcourt, R.L., Kilian, A., Wilson, K.J. and Keese, K. (2002) Microbial B-Glucoronidase genes, gene products and uses thereof. United States Patent number 6, 391,547.

Jin, S.G., Komari, T., Gordon, M.P. and Nester, E.W. (1987) Genes responsible for the supervirulence phenotype of Agrobacterium tumefaciens A281. Journal of Bacteriology 169: 4417–4425.

King, Z.R., Bray, A.L., LaFayette, P.R. and Parrott, W.A. (2014) Biolistic transformation of elite genotypes of switchgrass (Panicum virgatum L.). Plant Cell Reports 33: 313–322.

Kondo, T., Hasegawa, H. and Suzuki, M. (2000) Transformation and regeneration of garlic (Allium sativum L.) by Agrobacterium-mediated gene transfer. Plant Cell Reports 19: 989–993.

Kvitko, B.H. and Collmer, A. (2011) Construction of Pseudomonas syringae pv. tomato DC3000 mutant and polymutant strains. In: Plant Immunity pp. 109–128. Springer.

Larsen, J.S. and Curtis, W.R. (2012) RNA viral vectors for improved Agrobacterium-mediated transient expression of heterologous proteins in Nicotiana benthamiana cell suspensions and hairy roots. BMC Biotechnology 12: 21.

Lazo, G.R., Stein, P.A. and Ludwig, R.A. (1991) A DNA transformation–competent Arabidopsis genomic library in Agrobacterium. Bio/technology 9: 963–967.

Leifert, C., Morris, C.E. and Waites, W.M. (1994) Ecology of Microbial Saprophytes and Pathogens in Tissue Culture and Field-Grown Plants: Reasons for Contamination Problems In Vitro. Critical Reviews in Plant Sciences 13: 139–183.

Lin, C.-Y., Donohoe, B.S., Ahuja, N., Garrity, D.M., Qu, R., Tucker, M.P., Himmel, M.E. and Wei, H. (2017) Evaluation of parameters affecting switchgrass tissue culture: toward a consolidated procedure for Agrobacterium-mediated transformation of switchgrass (Panicum virgatum). Plant Methods 13: 113.

Lippincott, B.B. and Lippincott, J.A. (1966) Characteristics of Agrobacterium tumefaciens auxotrophic mutant infectivity. Journal of Bacteriology 92: 937–945.

Lippincott, J.A., Webb, J.H. and Lippincott, B.B. (1965) Auxotrophic mutation and infectivity of Agrobacterium tumefaciens. Journal of Bacteriology 90: 1155–1156.

Liu, Y., Miao, J., Traore, S., Kong, D., Liu, Y., Zhang, X., Nimchuk, Z.L., Liu, Z. and Zhao, B. (2016) SacB-SacR gene cassette as the negative selection marker to suppress Agrobacterium overgrowth in Agrobacterium-mediated plant transformation. Frontiers in Molecular Biosciences 3: 70.

Lowe, K., La Rota, M., Hoerster, G., Hastings, C., Wang, N., Chamberlin, M., Wu, E., Jones, T. and Gordon-Kamm, W. (2018) Rapid genotype “independent” Zea mays L. (maize) transformation via direct somatic embryogenesis. In Vitro Cellular & Developmental Biology - Plant 54: 240–252.

Lowe, K., Wu, E., Wang, N., Hoerster, G., Hastings, C., Cho, M.-J., Scelonge, C., Lenderts, B., Chamberlin, M., Cushatt, J., Wang, L., Ryan, L., Khan, T., Chow-Yiu, J., Hua, W., Yu, M., Banh, J., Bao, Z., Brink, K., Igo, E., Rudrappa, B., Shamseer, P.M., Bruce, W., Newman, L., Shen, B., Zheng, P., Bidney, D., Falco, C., Register, J., Zhao, Z.-Y., Xu, D., Jones, T. and Gordon-Kamm, W. (2016) Morphogenic Regulators Baby boom and Wuschel Improve Monocot Transformation. The Plant Cell 28: 1998–2015.

Mann, D.G.J., King, Z.R., Liu, W., Joyce, B.L., Percifield, R.J., Hawkins, J.S., LaFayette, P.R., Artelt, B.J., Burris, J.N. and Mazarei, M. (2011) Switchgrass (Panicum virgatum L.) polyubiquitin gene (PvUbi1 and PvUbi2) promoters for use in plant transformation. BMC Biotechnology 11: 1–14.

Meng, Q., Liu, Z., Zhang, Y., Liu, C., Ren, F. and Feng, H. (2014) Effects of antibiotics on in vitro-cultured cotyledons. In Vitro Cellular & Developmental Biology - Plant 50: 436–441.

Murashige, T. and Skoog, F. (1962) A revised medium for rapid growth and bio assays with tobacco tissue cultures. Physiologia Plantarum 15: 473–497.

Nester, E.W. (2015) Agrobacterium: nature’s genetic engineer. Frontiers in Plant Science 5: 730.

Ogawa, Y., Shirakawa, M., Koumoto, Y., Honda, M., Asami, Y., Kondo, Y. and Hara-Nishimura, I. (2014) A simple and reliable multi-gene transformation method for switchgrass. Plant Cell Reports 33: 1161–1172.

Ondzighi-Assoume, C.A., Willis, J.D., Ouma, W.K., Allen, S.M., King, Z., Parrott, W.A., Liu, W., Burris, J.N., Lenaghan, S.C. and Stewart, C.N. (2019) Embryogenic cell suspensions for high-capacity genetic transformation and regeneration of switchgrass (Panicum virgatum L.). Biotechnology for Biofuels 12: 290.

R Core Team (2021) R: A Language and Environment for Statistical Computing. R Foundation for Statistical Computing, Vienna, Austria. https://www.R-project.org/

Ranch, J.P., Liebergesell, M., Garnaat, C.W. and Huffman, G.A. (2010) Auxotrophic Agrobacterium for plant transformation and methods thereof. United States Patent number 8, 334, 429.

Rotem, O., Biran, D. and Ron, E.Z. (2013) Methionine biosynthesis in Agrobacterium tumefaciens: study of the first enzyme. Research in Microbiology 164: 12–16.

Schneider, C.A., Rasband, W.S. and Eliceiri, K.W. (2012) NIH Image to ImageJ: 25 years of image analysis. Nature Methods 9: 671–675.

Shackelford, N.J. and Chlan, C.A. (1996) Identification of antibiotics that are effective in eliminating Agrobacterium tumefaciens. Plant Molecular Biology Reporter 14: 50–57.

Somleva, M.N., Tomaszewski, Z. and Conger, B.V. (2002) Agrobacterium-mediated genetic transformation of switchgrass. Crop Science 42: 2080–2087.

Stewart, C.N., Jr. and Via, L.E. (1993) A rapid CTAB DNA isolation technique useful for RAPD fingerprinting and other PCR applications. BioTechniques 14: 748–750.

Wu, H.-Y., Liu, K.-H., Wang, Y.-C., Wu, J.-F., Chiu, W.-L., Chen, C.-Y., Wu, S.-H., Sheen, J. and Lai, E.-M. (2014) AGROBEST: an efficient Agrobacterium-mediated transient expression method for versatile gene function analyses in Arabidopsis seedlings. Plant Methods 10: 1–16.

Xi, Y., Fu, C., Ge, Y., Nandakumar, R., Hisano, H., Bouton, J. and Wang, Z.-Y. (2009) Agrobacterium-mediated transformation of switchgrass and inheritance of the transgenes. Bioenergy Research 2: 275–283.

Zhi, L., TeRonde, S., Meyer, S., Arling, M.L., Register Iii, J.C., Zhao, Z.-Y., Jones, T.J. and Anand, A. (2015) Effect of Agrobacterium strain and plasmid copy number on transformation frequency, event quality and usable event quality in an elite maize cultivar. Plant Cell Reports 34: 745–754.

